# A simple inland culture system provides insights into ascidian post-embryonic developmental physiology

**DOI:** 10.1101/2024.08.16.608202

**Authors:** Birthe Thuesen Mathiesen, Mayu Ohta, Boris Pinto De Magalhaes, Chiara Castelletti, Vincenzo Perria, Lionel Christiaen, Naoyuki Ohta

**Affiliations:** Michael Sars Centre, University of Bergen, Bergen, Norway; Center for Developmental Genetics, Department of Biology, New York University, New York, NY, USA

## Abstract

Maintenance and breeding of experimental organisms are fundamental to life sciences, but both initial and running costs, and hands-on zootechnical demands can be challenging for many laboratories. Here, we aimed to further develop a simple protocol for reliable inland culture of tunicate model species of the *Ciona* genus. We cultured both *Ciona robusta* and *Ciona intestinalis* in controlled experimental conditions, with a focus on dietary variables, and quantified growth and maturation parameters. From statistical analysis of these standardized datasets, we gained insights into the post-embryonic developmental physiology of *Ciona*, and inferred an improved diet and culturing conditions for sexual maturation. We showed that body length is a critical determinant of both somatic and sexual maturation, which suggests the existence of systemic control mechanisms of resource allocation toward somatic growth or maturation, and supports applying size selection as a predictor of reproductive fitness in our inland culture, to keep the healthiest animals at low density in the system. In the end, we successfully established a new protocol, including size selection, to promote both sperm and eggs production. Our protocol using small tanks will empower researchers to initiate inland *Ciona* cultures with low costs and reduced space constraints.

## Introduction

Popular model organisms for experimental research are typically broadly available across laboratories, benefitting from straightforward and inexpensive culture systems, lest difficulties will be imparted onto the projects (Matthews and Vosshall 2020; Bertile *et al*. 2023). Thus, it is essential to culture and maintain experimental organisms in the laboratory. However, this can represent substantial costs, limiting the accessibility of experimental models. Therefore, lowering initial and running costs for model organisms is poised to broaden their usability for experimental research.

*Ciona robusta* (also known as *Ciona intestinalis* type A) and *Ciona intestinalis* have emerged as chordate model organisms for molecular developmental and cell biology, as well as genomics, evolutionary biology, ecology and toxicology. Their phylogenetic position as the sister group to vertebrates in the chordate phylum, and the stereotypical and simple development of their relatively transparent embryo containing a compact genome, without the whole genome duplication observed in vertebrates, have propelled them as popular models for a wide range of studies (Satoh 2003; Lemaire 2011; Stolfi and Christiaen 2012). *Ciona* species are distributed worldwide and *Ciona robusta* is still invading new ecosystems (Lambert and Lambert 1998; de Oliveira Marins *et al*. 2009; Brunetti *et al*. 2015; Wilson *et al*. 2022), thus also becoming a model animal for applied ecology, especially for their invading of non-native environment by global shipping and global warming.

The first draft of the whole genome sequence was published in 2002 (Dehal *et al*. 2002), alongside fundamental datasets for molecular genomics research such as EST sequencing (Satou *et al*. 2001b, 2002a) and cDNA library (Satou *et al*. 2002b). Community databases for *Ciona* community (Satou *et al*. 2005, 2022; Hotta *et al*. 2007, 2020; Tassy *et al*. 2010; Endo *et al*. 2011; Dardaillon *et al*. 2020) have empowered progress in various scientific fields including genetics (Veeman *et al*. 2011), genomics (Keshavan *et al*. 2010; Hendrix *et al*. 2010; Kubo *et al*. 2010; Satou *et al*. 2015, 2021; Racioppi *et al*. 2019), and developmental and cell biology(Inaba *et al*. 2007; Lemaire *et al*. 2008; Lemaire 2009; Satou and Imai 2015; Satoh 2016; Oda-Ishii and Satou 2018). Molecular perturbation based on electroporation and microinjection with endonucleases (Kawai *et al*. 2012; Treen *et al*. 2014; Sasaki *et al*. 2014; Stolfi *et al*. 2014) and with morpholino antisense oligonucleotides(Satou *et al*. 2001a; Imai *et al*. 2006; Christiaen *et al*. 2009a; Ohta and Satou 2013; Waki *et al*. 2015) have been established for functional analysis through reverse genetics. Even though much progress has been made, these loss-of-function (LOF) phenotypes are typically induced transiently in wild-caught animals, following *in vitro* fertilization and short-term cultures in Petri dishes, rendering the analysis of late/post-embryonic stages more challenging. This also complicates the analysis with uncontrolled biological and technical variability, such as mosaicism, variable efficacy and penetrance, and uncharted artifacts and off-target effects.

To overcome the disadvantage of transient LOF, forward genetics strategies have been implemented (Jiang *et al*. 2005; Veeman *et al*. 2008; Chiba *et al*. 2009) and stable transgenic lines were established for reporter constructs and both forward and reverse genetics (Sasakura *et al*. 2005, 2012). Stable transgenesis and germline mutagenesis have been established in *Ciona* by using the Tc1/mariner family transposon *Minos* (Sasakura *et al*. 2003; Matsuoka *et al*. 2005; Hozumi *et al*. 2010; Horie *et al*. 2011; Yoshida and Sasakura 2012; Sasakura 2018). However, these genetically modified organisms can only be developed and maintained in closed culture systems that prevent escape to nature (Joly *et al*. 2007).

Several culture facilities have been developed, including open systems directly connected to the ocean, thus providing unlimited flow of seawater and planktonic food for ascidians (Hendrickson *et al*. 2004; Joly *et al*. 2007; Veeman *et al*. 2011). On the other hand, inland closed culture systems need to dedicated care and, whether it is automated or not, run at higher costs (Henry *et al*. 2020), especially to maintain a balance of water quality and food availability, a key factor in *Ciona* cultures (Joly *et al*. 2007; Zupo *et al*. 2020). Here, we used a principled science-based approach to test various culturing variables, including concentrations of microalgae, and monitored growth and maturation of the animals to further improve our culture protocol for *Ciona* species. This approach revealed several biological insights into the post-embryonic developmental physiology of *Ciona* and yielded a simple provisional protocol for animal culture.

## Materials and Methods

### Animal culture and observation

Wild-type animals of *Ciona robusta* (a.k.a *Ciona intestinalis* type A) were collected in California in the U.S.A, and shipped to New York. Then, eggs and sperm were surgically collected from 5-6 mature adults and were pooled on Petri dish for fertilization. The fertilized eggs were transferred to Petri dishes with artificial sea water (ASW; Bio actif sea salt, Tropic Marin), and 50-100 larvae were transferred into new Petri dishes on the next day to let them proceed metamorphosis like as previously reported(Swalla 2004; Joly *et al*. 2007). The salinity of the sea water was set at 32-34 ppt. The juveniles were shipped from New York in the U.S.A to Bergen, Norway. Wild-type animals of *Ciona intestinalis* were collected in Bergen, Norway. Eggs and sperm were surgically collected from 5-6 mature adults in each fertilization. The fertilized eggs were cultured on Petri dishes in ASW, and larvae were transferred to new Petri dishes on the next day. *Ciona robusta* and *Ciona intestinalis* juveniles were cultured at 18 ℃ and 14 ℃, respectively. ASW was refreshed, and the animals were fed every Monday, Wednesday and Friday. The Petri dish was filled with 20mL of ASW. The 0.8L and 10L tanks (Plastic storage box 0.8 L, Coline, 44-1740-1, Clas Ohlson; Round storage containers 12 qt, RFSCW12, Cambro) were filled with 0.7 L and 7 L of ASW, respectively. The Petri dishes were put in incubators (Model KBW 720, Growth chambers with light, Binder), which were set 14 ℃ and 18 ℃, and the 0.8 L and 10 L tanks were kept in our facility at 18 ℃ room temperature. Usually, *Ciona* juveniles have 0.5 mm body length at day 5-6 before they start to be fed, so we considered their initial body length as 0.5 mm. The body length of the animals was measured by a ruler under microscope (SZX10, Olympus; SMZ1270, Nikon). Images were taken by camera (HDMI16MDPX, DeltaPix) set up on the microscope (SMZ1270, Nikon).

### Algae culture

We used the diatom *Chaetoceros calcitrans* (CCAP 1010/11 *Chaetoceros calcitrans* fo. Pumilus, Culture Collection of algae & protozoa) and the haptophyte *Isochrysis sp*. (CCAP 927/1 *Iso* Culture Collection of algae & protozoa) which were cultured as previously described(Joly *et al*. 2007; Ohta *et al*. 2020). The concentration of algae in ASW was calculated by optical density (OD) at 600 nm using a spectrophotometer (Du-640, Beckman) as previously reported (Bouquet *et al*. 2009). Then, we prepared an initial condition (C1) with a concentration of 4.0 x 10^5^ cells/mL of each alga. And then prepared the other conditions like C3 and C10, that have 3 and 10 times cells from C1, respectively.

### Food supplement

We used an Oyster feast (Reef Nutrition, California in the U.S.A) as a food supplement to feed it to *Ciona* young adults in addition to regular algae. One drop of the Oyster feast was used for 3 L of ASW. The food supplement was used after animals had maturated enough to have orange pigment organ (OPO) at the tip of sperm duct in *Ciona robusta* animals.

### Plasmid DNA construction

Cr-Ef1α>msfGFP plasmid was made as below; monomeric-super-folded GFP was isolated from pC034-LwCas13a-msfGFP-2A-Blast plasmid (Addgene, #91924; (Abudayyeh *et al*. 2017)) by PCR using NotI_msfgfp_for (5’-AAAGCGGCCGCAACCATGGTGAGTAAAGGTGAAGAAC-3’) and EcoRI_msfgfp_rev (5’-CTGGAATTCTCACTTGTACAGCTCATCCATAC-3) primers. The PCR product was digested with NotI (NEB, R3189L) and EcoRI (NEB, R3101S), and was subcloned into *Ef1a* promoter that was obtained by digesting with NotI and EcoRI from Cr-Ef1α>Cas9(Stolfi *et al*. 2014). These 2 DNA fragments were ligated with T4 DNA ligase (Promega, M1804).

### Electroporation with the eggs from inland-cultured animals

We dissected 5 and 3 animals of *Ciona robusta* raised in C1 and C1+Oyster conditions as mixture to get sperm and eggs at 90 days post fertilization (dpf) at the 3^rd^ round of culture. And, we dissected in total 58 animals of *Ciona intestinalis* raised in C1.5 and C1.5+Oyster as mixture at 111, 120 and 133 dpf at the 4^th^ round of culture. The eggs were dechorionated and fertilized to be used for electroporation, which was described previously(Christiaen *et al*. 2009b). Plasmid DNAs were amplified by NucleoBond Xtra Midi (Macherey Nagel, 740410.100) and used for electroporation.

The detailed protocol was described previously(Christiaen *et al*. 2009b). Forty μg of Cr-Mesp>hCD4::GFP and 20 μg of Cr-Ef1α>msfGFP were used for electroporation in 700 μL scale of electroporation on the electroporation system (BTX, Gemini twin wave electroporation system). Exponential decay wave protocol on 50 voltage was used. The electroporated eggs were cultured at 14 ℃. These embryos were fixed at Stage 13 and Stage 23(Hotta *et al*. 2007) by MEM-FA (4% Formaldehyde, 0.1M MOPS, 0.5M NaCl, 1mM EGTA, 2mM MgSO4). We used Chicken anti-gfp (Abes Labs, GFP-1020) and anti-chicken-alexa488 (Fisher scientific, #10286672) antibodies to enhance the signals from Cr-Mesp>hCD4::GFP by immunostaining(Ohta and Satou 2013; Ohta and Christiaen 2024). The images were taken by FV3000 confocal microscope (Olympus) and were analyzed with Cell Sens (Olympus).

### Statistical analysis

We used Microsoft Office Excel to score and summarize our data, and analyzed the data by R in RStudio. The details of statistics were described in Table 1. The 1^st^ and 4^th^ round of *Ciona intestinalis* culture in Figure 2-3, 6 and S1 had 2 biological replications. The 2^nd^ and 3^rd^ round of *Ciona robusta* culture in Figure 2-6 and S2-3 had 3 and 2 biological replications, and 5 technical replications.

**Table 1.** Summary of cultures.

| round | species | aim | start |  | after selection |  | matured |  |  | data points | replicates |
| --- | --- | --- | --- | --- | --- | --- | --- | --- | --- | --- | --- |
|  |  |  | animal | Petri dish | animal | Petri dish | total | sperm + | egg + |  |  |
| 1st | *Ciona intestinalis* | test food concentration | 349 | 40 | N.D. | N.D. | N.D. | N.D. | N.D. | 2094 | 2 |
| 2nd | *Ciona robusta* | test animal concentration | 675 | 57 | 125 | 40 | 99 | 21 | 4 | 3341 | 3 |
| 3rd | *Ciona robusta* | test new protocol | 490 | 63 | 128 | 61 | 103 | 76 | 35 | 909 | 2 |
| 4th | *Ciona intestinalis* | test new protocol | N.D. | N.D. | 160 | 80 | 140 | 136 | 76 | 794 | 2 |

### Data Availability Section

This study includes no data deposited in external repositories.

## Results

### Monitoring growth and maturation to optimize inland culture of *Ciona*

To expand usage of *Ciona robusta* and *Ciona intestinalis* as model organisms, especially in laboratories located far from the Pacific Ocean, we aimed to further develop a stable inland culture system with lower costs, smaller space and reduced maintenance. First, to control and establish a reference for food quantity, we measured the number of algae particles from the optical density (OD) of cultures, as previously reported for *Oikopleura* inland cultures (Bouquet *et al*. 2009). Based on previous reports of successful inland culture (Malfant *et al*. 2017), we set condition 1 (C1) to contain 4.0 x 10^5^ cells/mL of each alga as initial standard. To assess the impact of food concentration, we also used conditions such as C2 and C3, which had 2 and 3 times more algae than C1, respectively. To monitor growth and somatic maturation through the juvenile period, we measured body length, counted the numbers of gill slits and observed atrial siphon fusion (Figure 1A-B) (Chiba *et al*. 2004; Ohta *et al*. 2010; Sasakura and Hozumi 2018). We scored the presence of the orange pigment organ (OPO) at the tip of sperm duct as a species-specific sign of somatic maturation in *Ciona robusta*, as well as an early sign of sexual maturation, in addition to presence of sperm and eggs (Figure 1C-D). Previous reports indicated that *Ciona* animals can be cultured in Petri dishes at juvenile stages, and should be transferred to larger culture systems when they reach a body length of 2 mm (Joly *et al*. 2007). Here, we introduced a 0.8 L tank for small scale culture, which allowed us to keep animals in smaller spaces at reduced initial cost (Figure 1E). We transferred juvenile-containing Petri dishes into larger tanks once they completed atrial siphon fusion to reach the 2^nd^ juvenile stage and had grown over 2 mm in body length (Chiba *et al*. 2004; Joly *et al*. 2007; Ohta *et al*. 2010). We monitored sexual maturity over a time course once animals had been moved into either 0.8 L or 10 L tanks (Figure 1F). From our first and second rounds of observation in both *Ciona intestinalis* and *robusta*, we collected 2,094 and 3,341 data points from an initial population of 349 and 675 individuals distributed in 40 and 57 Petri dishes, respectively (Table 1). In the following sections, we analyze this dataset to understand and improve the conditions for growth, and somatic and sexual maturation.

**Figure 1.**
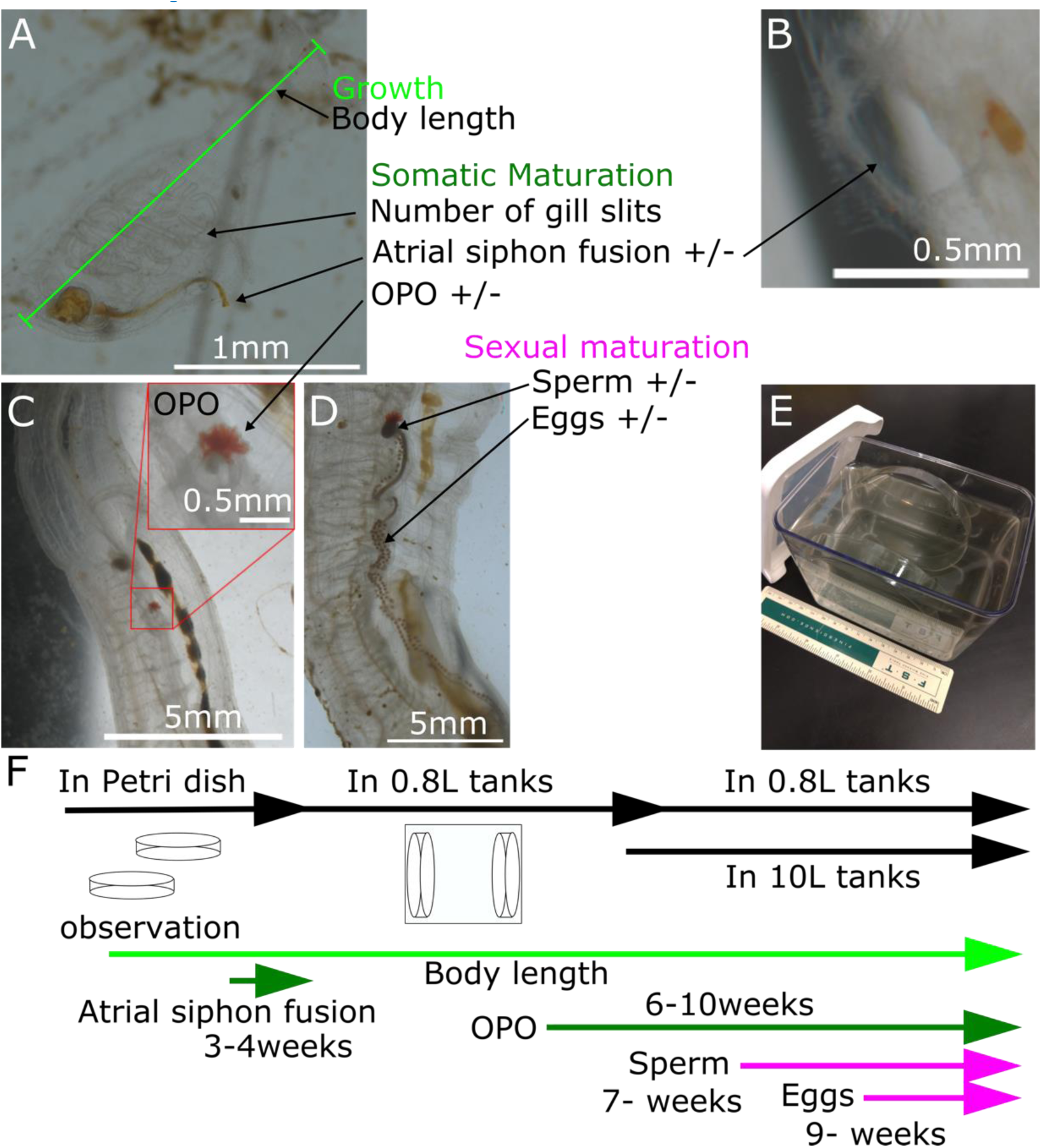
Monitoring growth and maturation of *Ciona robusta* and *Ciona intestinalis* in inland culture. (A) Body length was measured as a parameter for growth, and the number of gill slits was counted as a parameter for somatic maturation. Atrial siphon fusion was observed as a parameter for somatic maturation in juvenile stage as binary data. (B) We scored the completion of atrial siphon fusion when the two openings become one. (C) Orange pigment organ (OPO) was observed as a parameter for somatic maturation in adult stage as binary data. (D) Sperm and eggs were observed as parameters for sexual maturation as binary data. (E) The 0.8L tank was used for inland culture. (F) Schedule for culturing and observation.

### An optimal range of food availability for somatic growth

In our 1^st^ round of culture and monitoring with locally caught *Ciona intestinalis*, we aimed to test the impact of food concentration on growth in a Petri dish. We prepared 6 conditions from condition 0 (C0), that had no algae, to condition 30 (C30), that had 30 times more algae particles than the C1 (Figure S1A-F). Animals placed in the C30 condition survived at significantly lower rate than those placed in other conditions, including the C0 condition, which survived at a higher rate until day 26 (Figure S1G). Juveniles in the C30 condition failed to grow and died due to an excess of algae that accumulated in the Petri dishes, and polluted the seawater, causing substantial increase in pH, which correlated negatively with survival (Figure 1F and S1I). The fact that juveniles in the C0 condition survived at rates equivalent to fed animals, and even grew in the absence of food, revealing the existence of mechanisms allowing survival and even development despite starvation (Ohta et al., in preparation).

Next, we leveraged our longitudinal monitoring of individual animals to study the impact of food availability on somatic growth. Compared to the C1 reference, animals cultured in the C0.1 condition displayed shorter body length and lower growth rates during the first 26 days, confirming the intuitive expectation that food availability can be limiting during the first month (Figure 2A-B and S1H). On the other hand, animals cultured in the C10 condition also exhibited lower growth rates and 26-day length compared to C1 and C3 animals, which thus define an “optional range” of algae concentration (Figure 2A-B and S1). Thus, we mainly used the C1 to C3 in our 2^nd^ round of culture and monitoring with *Ciona robusta*.

**Figure 2.**
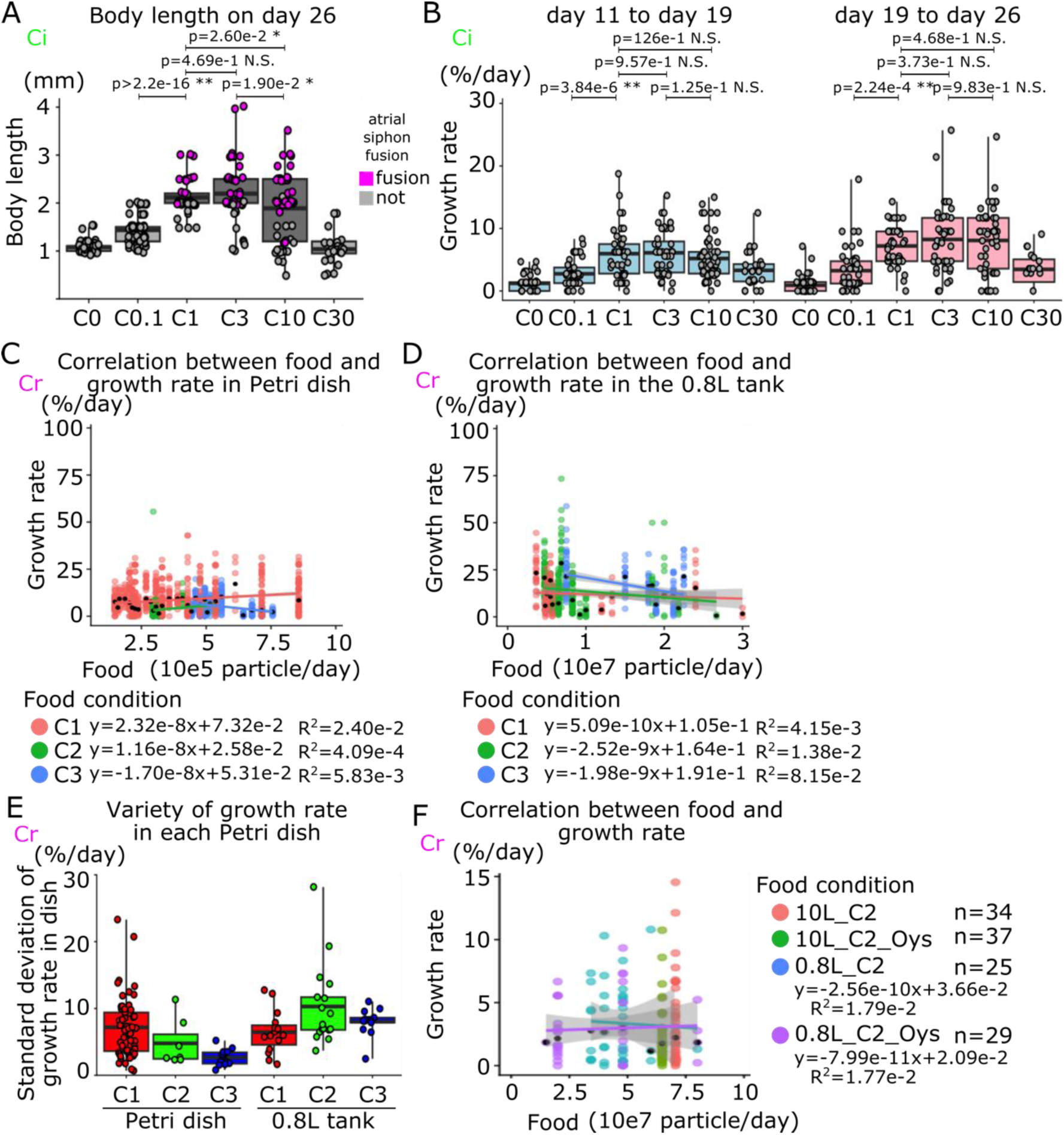
Effect of food availability on growth. (A) Box and dot plots show the body length of juveniles on day 26 in each dietary condition. (B) Box and dot plots show the growth rate of individual juveniles before and after day 19 in each dietary condition. (C) Correlation between food availability and growth rate in Petri dishes was shown in scatter plot. Each colored line indicates a fitting curve. This plot uses a part of the data of food particle between 10e5-e6 particle/day. (D) Correlation between food availability and growth rate in a 0.8L tank is shown in scatter plot. Each colored line shows fitting curve. This plot uses a part of data on food particle between 10e6-30e7 particle/day. (E) Variability of body length of each juvenile in culture system is shown as standard deviation of body length in each Petri dish. (F) Correlation between food accessibility and growth rate in each dietary condition after animal selection is shown in scatter plot. Each colored line shows fitting curve. Each dot shows individual juveniles observed at one time point on A-D, F. Black dots show median of the growth rate among each dietary condition on particle base or each condition of different animal numbers in a Petri dish on C-D, F. The color indicates dietary conditions of the Petri dishes on C-D, F. Magenta and gray show completion of atrial siphon fusion or not on A. Species of animals that are used in each graph are indicated by green “Ci” and magenta “Cr” for *Ciona intestinalis* and *Ciona robusta*, respectively. p-value was calculated by t-test on A and B. p>0.05; N.S, 0.05>p>0.01; *, 0.01>p; **.

Leveraging our controlled diet, we estimated the number of algae particles accessible to each individual juvenile per day, which allowed us to plot the relationship between food availability and growth for animals cultured in inland culture system (Figure 2C-D). Notably, individual growth rates were highly variable, even between animals cultured in the same Petri dish and tank (Figure 2E). Nevertheless, statistical analyses based on large numbers of data points indicated that animal growth did not correlate with food availability, suggesting that, with the “optimal range” of food availability, stochastic variation and/or other parameters, such a genetic polymorphism or initial egg composition, impact individual growth rates more than minor differences in food availability. Because confluency of animals in a Petri dish is related to the number of food particles that each juvenile can access, we expected animal density in a Petri dish to impact somatic growth, as previously suggested from qualitative observations (Joly *et al*. 2007). However, our longitudinal data failed to capture a correlation between growth rate and food availability for animals cultured in the permissive range (Figure S2A-B).

To further explore the correlation between food availability and somatic maturation, some of the animals were selected from the population based on their body length and the presence of an orange pigment organ (OPO), yielding 125 animals distributed in 40 Petri dishes in the 2^nd^ round of culture (Table 1). We monitored growth and maturation after transferring Petri dishes to 0.8 L or 10 L tanks, using the C2 condition, with or without “Oyster feast” supplement (Figure 2F, 4 S2 and S3). As for the first round of culture, individual growth rates did not correlate with calculated food availability (Figure 2F), further indicating that, within a permissive/optimal range, individual-to-individual growth variation is not explained by food availability.

### Somatic maturation correlates with food availability

Following attachment to a substrate, *Ciona* larvae metamorphose into juveniles, which acquire 2 pharyngeal gill slits and 2 atrial siphon primordia on either side. Animals then undergo a remarkable process of somatic maturation whereby atrial siphon primordia migrate and fuse dorsally, completing atrial siphon fusion by the end of the 1^st^ juvenile stage, until that point they were cultured in Petri dishes in a temperature-controlled incubator (Figure 1F).

In another round of culture, this time with *Ciona intestinalis* animals collected from the Norwegian shores, we varied dietary conditions and longitudinally monitored pharyngeal gill slit formation and atrial siphon fusion in individual animals (Figure 3A-C). While atrial siphon fusion is a progressive process, we scored it as a simplified binary event to analyze the impact of dietary conditions here. The percentage of atrial siphon fusion thus represents the number of juveniles that have completed the process. As observed for *Ciona robusta*, the C1, C3 and C10 conditions defined an optimal range for growth, but also somatic maturation as assayed by gill slit formation and atrial siphon fusion. Indeed, the C3 condition maximized both parameters in *Ciona intestinalis*, where gill slit numbers were highly correlated with animal length (Figure 3A-C). On the other hand, there were no significant differences in rates of atrial siphon fusion between optimal dietary conditions. Notably, while animals in C1 and C3 conditions displayed similar individual growth rate, C3 animals appeared to mature more rapidly, suggesting that beyond a growth optimal, additional resources are invested into somatic maturation. Therefore, we considered that the C3 condition, which showed the best growth and maturation, is the optimal dietary condition for inland culture of *Ciona intestinalis* at 14 ℃ with 8 juveniles in a Petri dish.

**Figure 3.**
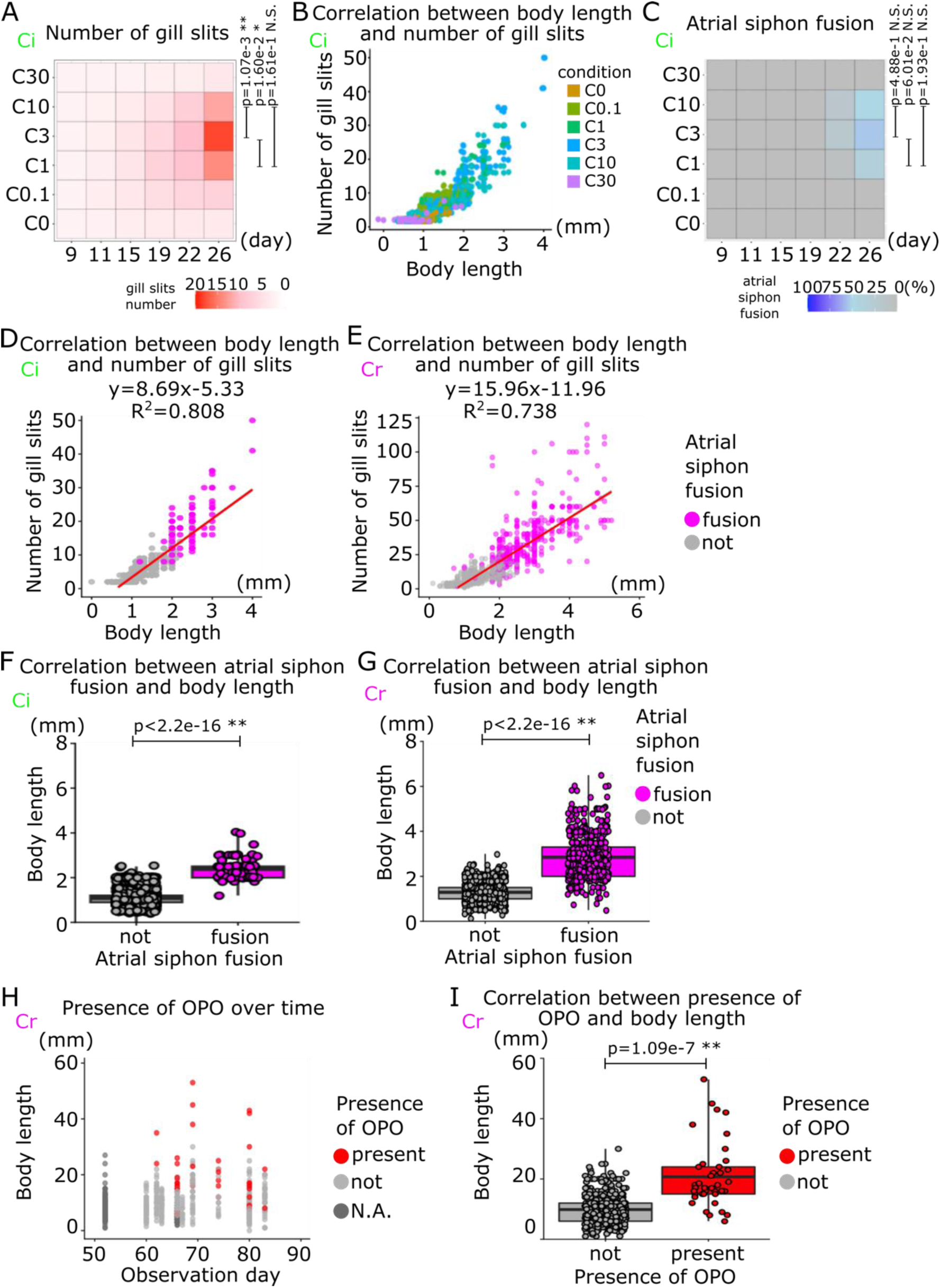
Correlation between food availability and somatic maturation. (A) Average of the numbers of gill slits on half side of juveniles over time in each dietary condition is shown in heat map. (B) Correlation between body length and number of gill slits of each dietary condition is shown in scatter plot. (C) The rate of juveniles which have done atrial siphon fusion over time in each dietary condition is shown in heat map. (D-E) Correlation between body length, the number of gill slits and atrial siphon fusion is shown in scatter plot. The red line indicates a fitting curve. (F-G) Box and dot plot show correlation between atrial siphon fusion and body length. (H) Body length and presence of orange pigment organ (OPO) over time are shown in scatter plot. (I) Box and dot plot show correlation between presence of OPO and body length. Each dot shows individual animals observed at one time point on B, D-I. Magenta and gray show completion of atrial siphon fusion or not on D-G. Red and gray show presence of OPO or not at the tip of sperm duct on H and I. Species of animals which are used in each graph are indicated by green “Ci” and magenta “Cr” for *Ciona intestinalis* and *Ciona robusta*, respectively. p-value was calculated by t-test on A, C, F, G and I. p>0.05; N.S, 0.05>p>0.01; *, 0.01>p; **.

To further fully understand growth and maturation of *Ciona* juveniles at their 1^st^ juvenile stage, we plotted the parameters of body length, gill slit and atrial siphon fusion together in dataset in the 1^st^ and 2^nd^ rounds of culture, and sought to extend our observations to *Ciona robusta* (Figure 3D-E). In both species, the number of gill slits was highly correlated with body length (R^2^ = 0.808 and R^2^ = 0.738). Of note, water temperature affects animal growth and maturation, and *Ciona robusta* and *intestinalis* have different preferences (Malfant *et al*. 2017); this led the body length and the number of gill slits to reach different values between species, but the strong correlation remained.

Atrial siphon fusion is a remarkable morphogenetic event, resulting in the merging of two pre-atrial siphons into a single atrial siphon opening after fusion. In both *Ciona* species, atrial siphon fusion was observed in animals with bodies longer than 2 mm (Figure 3F-G), further suggesting the existence of growth thresholds for animals to undergo somatic maturation.

Once *Ciona* juveniles completed atrial siphon fusion, they enter the 2^nd^ juvenile stage with one mature atrial siphon, and almost assume their definitive adult shape (Chiba *et al*. 2004). At the 2^nd^ juvenile stage, animals were cultured in 0.8 L tanks with 2 Petri dishes attached vertically at either side of the tank (Figure 1E), and we continued to monitor animal growth and maturation. As a species-specific sign of maturation, the orange pigment organ (OPO) becomes visible at the tip of sperm duct in *Ciona robusta* as a character of *Ciona robusta* (Millar 1953; Hoshino and Tokioka 1967; Caputi *et al*. 2007; Ohta *et al*. 2010, 2020; Sato *et al*. 2012, 2014; Tajima *et al*. 2020). We thus used OPO formation to monitor somatic maturation at the 2^nd^ juvenile stage in *Ciona robusta* (Figure 1C, 3G). As for gill slit formation and atrial siphon fusion, OPO formation correlated with animal length as animals with OPO were significantly larger (Figure 3H-I). Taken together, these results identified a range of food availability for optimal growth and somatic maturation in *Ciona* juveniles, revealing a correlation that suggests the existence of (a) gating mechanism(s) whereby, past certain size thresholds, nutritional resources no longer correlate with growth but instead promote somatic maturation.

### Sexual maturation in *Ciona robusta*

Having established defined culture conditions that optimize early growth and somatic maturation, we aimed to improve sexual maturation, which is essential to establish stable lines of genome-engineered animals. During the 2^nd^ round of culture with *Ciona robusta*, we monitored individual animals for sexual maturation in different dietary conditions; using 0.8 L and 10 L tanks culture systems, in the C2 condition supplemented or not with Oyster feast (Figure 4). After 118 days in culture, 21 and 4 animals had sperm and eggs, respectively, out of 99 survivors. Because *Ciona* animals produce sperm before eggs (Joly *et al*. 2007), egg-carrying animals also had sperm. Here, we tested 4 dietary conditions and monitored both growth, maturation, and sperm and egg production, but we did not detect statistically significant differences between the conditions on sperm and/or egg production (Figure 4A-B).

**Figure 4.**
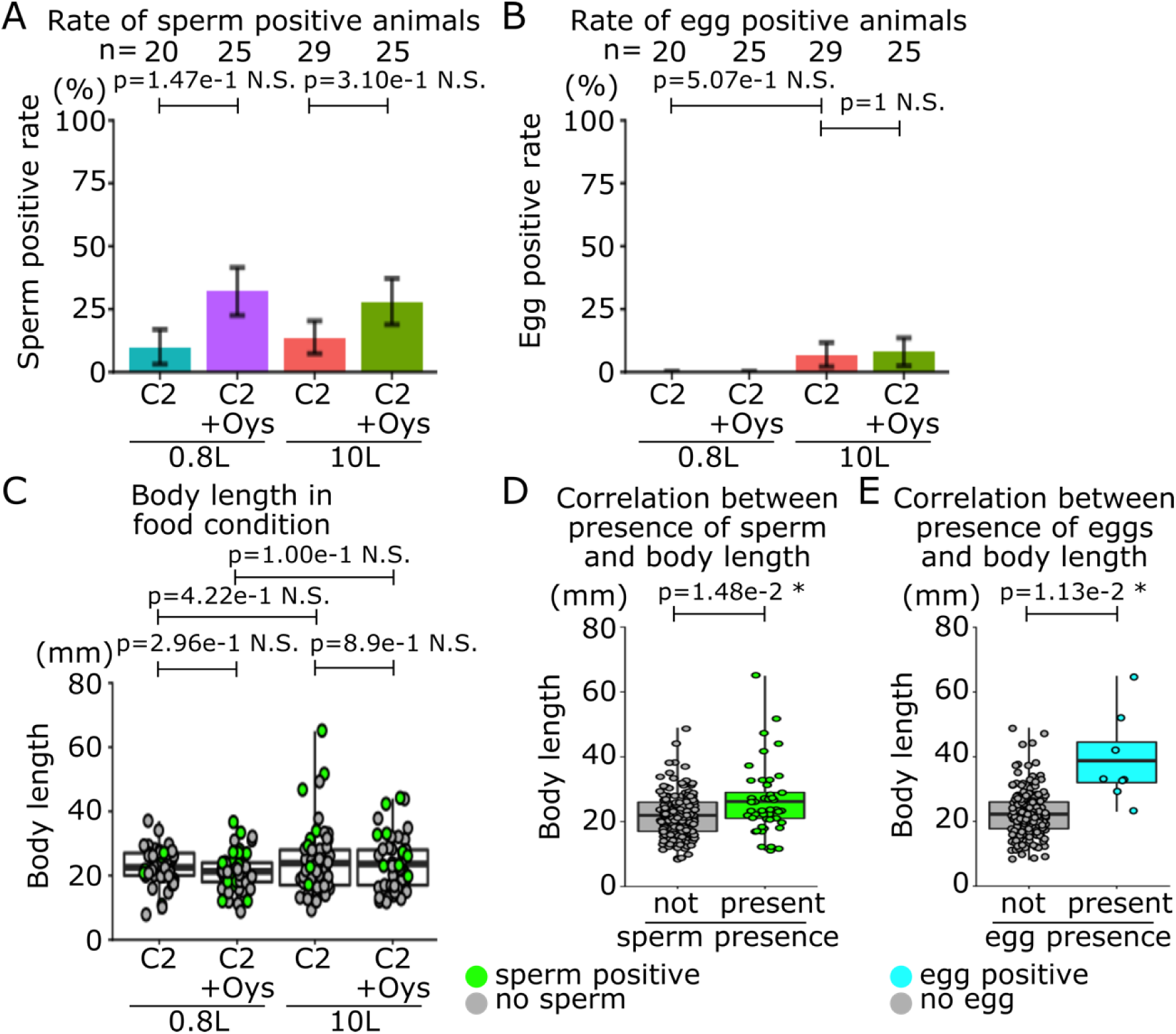
Sexual maturation of *Ciona robusta*. (A) Rate of sperm positive animals out of total animals in each dietary condition at the end of the culture are shown. (B) Rate of egg positive animals out of total animals in each dietary condition at the end of the culture are shown. (C) Box and dot plots show body length of animals in each dietary condition. (D) Box and dot plots show correlation between presence of sperm and body length. (E) Box and dot plots show correlation between presence of eggs and body length. Each dot shows individual animal observed at one time point on C-E. The color indicates dietary conditions of the animals in the 0.8L and 10L tanks with or without the Oyster feast as a supplemental food on A-B. Error bars show standard error on A-B. Green and gray show presence of sperm or not on C-D. Cyan and gray show presence of eggs or not on E. *Ciona robusta* is used for all the graphs in this figure. p-value was calculated by Fisheŕs exact test on A-B, and by t-test on C-E. p>0.05; N.S, 0.05>p>0.01; *, 0.01>p; **.

Consistent with data collected from the same animals at earlier stages (Figure 2F), there was no significant difference in body length of mature animals among the conditions (Figure 4C). Nevertheless, as observed for somatic maturation parameters, we detected a significant correlation between animal length and sexual maturation (Figure 4D-E). While smaller animals could still produce sperm, only the larger ones produced eggs, suggesting the existence of a size threshold on sexual maturation should be present in *Ciona* animal, as well as the size-gated threshold, which is more severe on egg production, explaining why it is typically easier to obtain cultured animals with sperm than eggs.

### Retrospective analysis of developmental trajectories for egg production

To understand how the animals reached an egg-producing state at the end of our observation, we extracted their corresponding data points from the whole dataset (Figure 5). Our data and analyses showed that somatic and sexual maturation correlated with animal length, indicating that monitoring animal growth should help understand of culture conditions on both growth and maturation (Figure 3 and 4). The population of egg-producing animals had larger body length over time than the population which produced neither sperm nor eggs at the end (Figure 5A). The growth rate history of the animals revealed a “growth burst” at around 1 month after fertilization (Figure 5B), consistent with a previous report (Joly *et al*. 2007). Notably, both populations experienced two growth bursts, while the egg-producing animals experienced it earlier. The timing of this growth burst, seemingly occurring after animals have completed somatic maturation, opens the intriguing possibility that, just as growth appears to gate somatic maturation at early juvenile stages, somatic maturation by early adult stages might be required for sexual maturation. Likely because the egg-producing animals had grown earlier than the others, they completed somatic maturation earlier (Figure 5C-E). This further indicated the existence of a size threshold for somatic maturation, and suggested that a somatic maturation gate determines sexual maturation, possibly in a time-dependent manner. In other words, the animals that grew too slowly failed to complete somatic maturation in the proper time window to produce eggs, at least in our experimental conditions.

**Figure 5.**
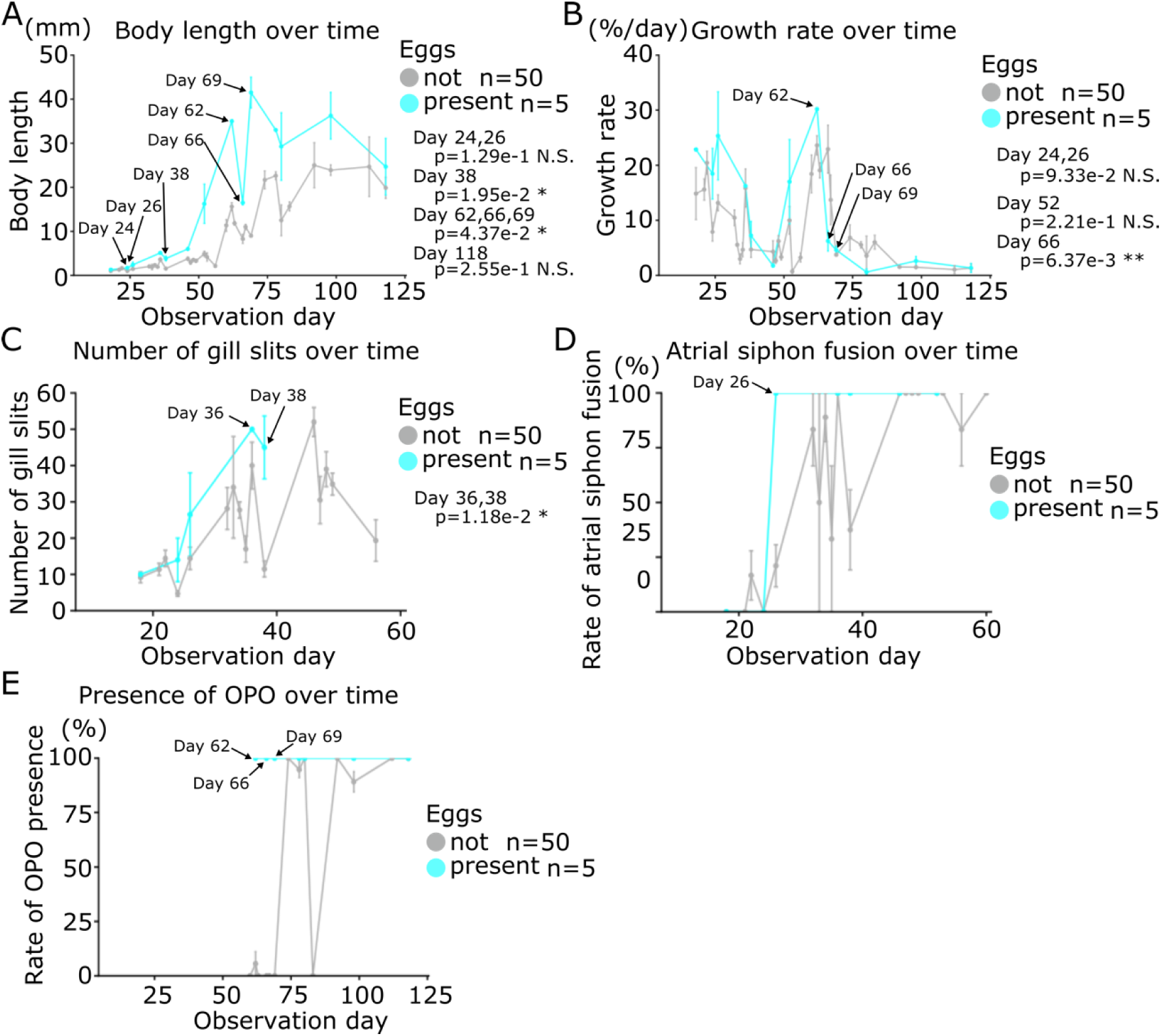
Trajectory of the egg-producing animals. (A-E) Trajectory of growth (A), growth rate (B), number of gill slits (C), atrial siphon fusion (D) and presence of orange pigment organ (OPO) (E) of the animals of egg-producing (cyan) and neither-sperm-nor-egg-producing (gray) during our culture are shown. *Ciona robusta* is used for all the graphs in this figure. p-value was calculated by t-test on A-B, D-E. p>0.05; N.S, 0.05>p>0.01; *, 0.01>p; **.

### A new protocol to promote egg production

To mimic the trajectory of growth and maturation that led to egg-producing animals, we established a new protocol based on the dietary conditions and the density of animals to maximize growth, and increase the number and proportions of egg-producing adults (Figure 6A). Even though juveniles fared better at low density in Petri dishes, we kept as many animals as possible to reduce the risk of animal loss at the 1^st^ juvenile stage. We thus started with 4 to 8 juveniles per Petri dish. To favor bigger animals, which were more likely to produce eggs (Figure 4E and 5), we introduced a size selection step by removing the smallest animals from Petri dishes before transfer into 0.8 L tanks. After that, we focused on either C1 or C2 conditions, with or without Oyster feast supplement, which allowed for animal growth and maturation. We set out to test this updated protocol and assay sperm and egg production through the 3^rd^ and 4^th^ rounds of culture of *Ciona robusta* and *intestinalis*, which we started with 63 and 80 Petri dishes (Table 1). The largest 128 juveniles were selected in Petri dishes before transfer into the 0.8 L tanks in the 3^rd^ round of *Ciona robusta* culture (Figure 6B). We monitored them longitudinally for 3 months when *Ciona* animals typically produce eggs (Joly *et al*. 2007). Similar to our 1^st^ and 2^nd^ rounds of culture, the largest animals also displayed more signs of somatic and sexual maturation than others (Figure S3). In the end, we obtained animals with both sperm and eggs (Figure 6C). In total, 73% (76/103) and 34% (35/103) animals produced sperm and eggs, respectively, out of 103 animals that survived until the end of observation in the 3^rd^ round of *Ciona robusta* culture, and 97% (136/140) and 54% (76/140) animals produced sperm and eggs, respectively, out of 140 animals that survived until the end of observation in the 4^th^ round of *Ciona intestinalis* culture (Figure 6, Table 1). These were improved figures compared to our 2^nd^ round of culture, where 21% (21/99) and 4% (4/99) animals had sperm and eggs, respectively, out of 99 survivors. In our 2^nd^ round of culture, we observed no significant difference between conditions with or without food supplement (Figure 4). By contrast, the Oyster feast supplement significantly improved sperm and egg production in the 3^rd^ culture, which used C1 as a base condition (Figure 6D-E).

**Figure 6.**
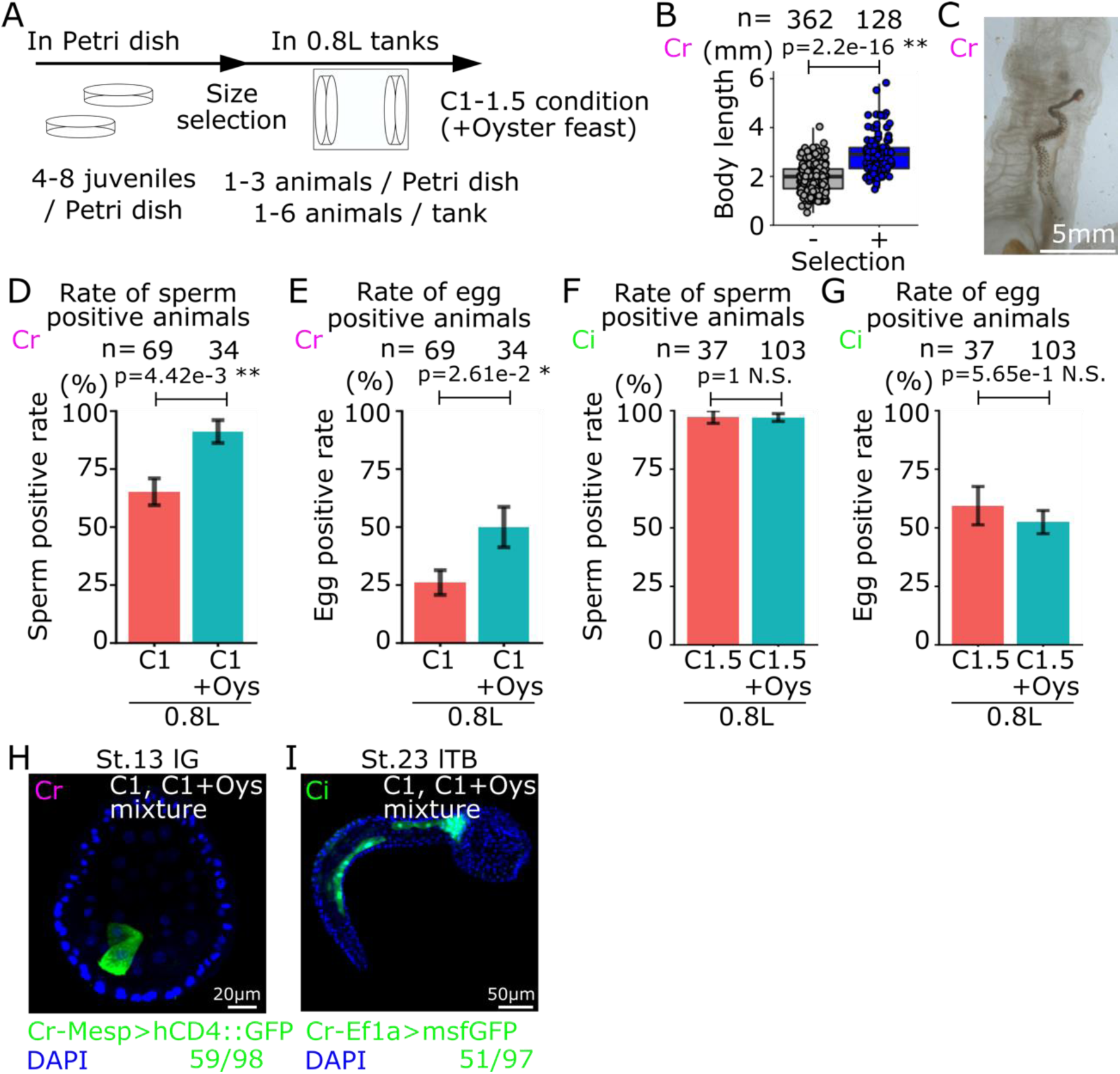
Verification of new protocol. (A) Schedule of culturing in our new protocol. In Petri dishes, 4 to 8 juveniles are cultured in C1 condition. After 3 weeks, the biggest 1-3 animals are selected in each Petri dish, then they are transferred into the 0.8L tank. At most 6 animals in the 0.8L tank are cultured in C1 condition with or without the Oyster feast. (B) The bigger juveniles in the population were selected by size selection. Blue and gray show selected and unselected juveniles, respectively. Each dot shows individual animal observed at size selection. (C) An image of the egg-producing animal raised in our inland culture system. (D-G) Rate of sperm (D and F) and egg (E and G) positive animals out of total animals in each dietary condition at the end of the culture with *Ciona robusta* (D-E) and *Ciona intestinalis* (F-G) are shown. (H-I) The eggs obtained from inland-cultured *Ciona robusta* (H) and *intestinalis* (I) were used for electroporation. Numbers show signal positive embryos out of total embryos. Error bars show standard error on D-G. Species of animals that are used in each graph and picture are indicated by green “Ci” and magenta “Cr” for *Ciona intestinalis* and *Ciona robusta*, respectively. p-value was calculated by t-test on B, and by Fisheŕs exact test on D-G. p>0.05; N.S, 0.05>p>0.01; *, 0.01>p; **.

Whereas there was no significant difference between conditions with or without food supplement in the 4^th^ round of culture, which used a C1.5 condition on *Ciona intestinalis*, and might suffice for sexual maturation (Figure 6F-G). These results suggested that additional food supplement can improve sperm and egg production, but the conflicting results do not clarify whether the supplements provided qualitative or quantitative benefits in these experiments.

Finally, since an essential goal of animal culture is to perform experimental manipulations, we used lab-produced eggs for electroporation to test if these were fit for experiments. We performed electroporation after egg dechorionation and fertilization, and obtained transgene-expressing embryos from both *Ciona robusta* and *intestinalis* adults that were raised our inland culture system in our 3^rd^ and 4^th^ rounds of culture (Figure 6H-I). This last result indicated that the eggs produced by animals raised in our inland culture system could be used for electroporation. Therefore, here we described an updated protocol to obtain fertile animals in an inland culture system with a reduced footprint and lower running costs.

## Discussion

Research using the Tunicate model *Ciona* still relies extensively on wild-caught animals, and thus remains impacted by seasonal variations and the inherent vagaries of climatic conditions. Mariculture can attenuate these limitations, and has been successfully used to produce mature wild-type animals (Sato *et al*. 2012, 2014). In Japan, the national bioresource project (NBRP; https://nbrp.jp/en/) has been providing wild-type adults of *Ciona robusta*, with a controlled genetic background (Sasakura *et al*. 2009), which were raised in the sea. Such mariculture system exploits natural resources in a cost-effective manner; however, it is not widely accessible and non-local and genetically modified organisms can obviously not be cultured in the sea. Circumventing this limitation, inland cultures of *Ciona* have been established world-wide, contributing to important research outputs. These ranged from transgenic animals made by transposon-mediated mutagenesis (Sasakura *et al*. 2003; Joly *et al*. 2007; Sasakura 2018) to development of an inbreed line (Satou *et al*. 2015), and multi-generation hybridization between closely related species (Ohta *et al*. 2020).

Nevertheless, inland culture systems remain suboptimal, costly and high maintenance. Here, we sought to develop a simplified yet effective culture system for the model species *Ciona robusta* and *Ciona intestinalis*. Focusing on such experimental variables as food quantity and quality, animal density and tank volume, we monitored individual animal growth, somatic and sexual maturation over 1- to 4-month time courses in a longitudinal manner. This principled approach and quantitative analyses allowed us to infer a simplified and improved protocol for animal culture from fertilized eggs to mature adults.

Beyond the practical advantages of a science-driven approach to zootechny, our results provide insights into post-embryonic developmental physiology, especially the impact of nutrition on growth, and somatic and sexual maturation. We first observed that, during the first month post-fertilization and metamorphosis, growth depends upon the availability of food, as expected, but a maximum growth rate is attained at relatively low concentration (C1) and additional resources (C3 or C10) do not translate into accelerated growth. On contrary, excess food (C30) was not entirely consumed by animals, decayed and caused detrimental pollution that reduced growth and survival. This pollution was marked by a significant increase in pH, which is known to affect *Ciona* development (Jones *et al*. 2023). While such trivial problem could be mitigated by using larger volumes, by more frequent water changes and by using complex recirculating systems, our data indicated that the corresponding costs and efforts are not warranted as 0-1-month-old animals did not grow faster above a low threshold of food availability.

On the other hand, our dataset and analysis revealed a significant correlation between animal size and somatic maturation, as indicated by gill slit numbers and atrial siphon fusion in 0.5 - 1 month old animals. Both gills slit formation and atrial siphon fusion are “dramatic” morphogenetic events that mark the transition from the 1^st^ to 2^nd^ juvenile stage (Chiba *et al*. 2004; Hotta *et al*. 2020). Our results indicate that, besides the observed correlation between animal size and somatic maturation, there appear to be a size threshold at ∼2 mm below which virtually no animal completed atrial siphon fusion. Remarkably, while intermediate food concentrations did not appear to augment somatic growth, increased food availability (C3 condition) seemed to accelerate somatic maturation. This suggests that, in the optimal growth regime, additional resources are allocated to somatic maturation rather than growth, even in relatively small animals (2 mm), which will later grow to larger sizes (up to 40 mm in our dataset). Of note, a first “growth burst”, characterized by high daily growth rates, coincides and/or slightly precedes the onset of somatic maturation at 20-25 dpf. Taken together, the growth plateau, size threshold for somatic maturation, differential resource allocations and growth burst suggest the existence of systemic mechanisms controlling the tempo of morphogenetic events during metamorphosis. It is tempting to speculate that diet-dependent endocrine systems control these transitions, as is the case in numerous other organisms.

During the last, 1 to 3/4 months, phase of our cultures, some adult animals reached sexually maturity, as evidence by the appearance of orange pigment organs (OPO) in *Ciona robusta*, as well as sperm and occasionally eggs in both species. As for somatic maturation, the likelihood of sexual maturation correlated with growth, and size thresholds appears to exist, albeit at higher values corresponding to approximately 20 and 30 mm for sperm and eggs, respectively (Figure 4D, lower limit of bounding box). Of note, a second growth burst occurred between 50 and 70 days, coinciding with the emergence of OPOs in *Ciona robusta*, and preceding the apparition of sperm and eggs. These observations suggest the existence of a second systemic control mechanism tying somatic growth and maturation with sexual maturation.

Of note, retrospective analysis indicated that animals that did not produce eggs also experienced a growth burst albeit with a delay, and were smaller on average (Figure 5A-B). This observation opens the intriguing possibility that a temporal gating mechanism operates in addition to a size threshold during the second systemic transition, although this remains to be confirmed and further analyzed.

Here, we attempted to convert these novel insights into the nutritional control of post-embryonic developmental physiology in *Ciona* into practical solutions for an improved and simplified inland culturing system. We sought to maximize food availability while reducing the risks of waste-induced pollution by focusing on the lowest effective concentration (C1-C1.5) while minimizing animal density, by curtailing per dish populations. Upon transfer of Petri dishes to 0.8 L tanks, we implemented a size-selection step to further control animal density and favor the largest animals, which fared better in prior cultures. We proposed that the growth and maturation trajectory of egg-producing animals provides an augmented timetable from juvenile to mature adult stage of *Ciona*, and builds upon previous detailed developmental tables until juvenile stages (Hotta *et al*. 2007, 2020).

This provisional protocol improved the outcome of this last round of culture with regards to sexual maturation with approximately 90% of the *Ciona robusta* animals producing sperm, and up to 50% producing eggs, compared to 25 and 10%, respectively, with as little as C1 vs. C2 supplemented with Oyster feast, and in 0.8 L vs. 10 L tanks (compared Figures 4 and 6). When using C1 however, the Oyster feast supplement appeared to increase the proportions of *C. robusta* animals producing sperm and eggs, compared to C2 (Figure 4) or *Ciona intestinalis* fed with C1.5 (Figure 5), suggesting that Oyster feed adds quantitatively and/or qualitatively to the animal diet required for sexual maturation. Almost 100% and 50% of the *Ciona intestinalis* animals produced sperm and eggs, respectively; irrespective of the use of Oyster supplement, suggesting that the latter did not add qualitative value to the C1.5 condition, which may already have been enough to sustain growth and maturation of the top animals.

In this study, we focused on food concentration and animal density in a small-scale inland culture system that supported sufficient animal growth and maturation to obtain gametes for experimental embryology, including a proof-of-concept electroporation. However, our curtailing approach and the small volumes used limit the quantity of animals and eggs that could be obtained, which remains limiting for larger scale experiments. We anticipate that several other parameters, which we did not test, could contribute to further optimizing this provisional protocol.

Regarding the diet and feeding regime, the new protocol is based on feeding and cleaning tanks on Mondays, Wednesdays and Fridays; but we do not know how quickly the animals “clear” the water, which might contrast with their ability to feed constantly on a steady concentration of microalgae in their natural habitat. *Ciona robusta* might be more sensitive to this “intermittent fasting” as it appeared to feed more intensely than *Ciona intestinalis* (Hoxha *et al*. 2018). As we saw, simply increasing the initial concentration of food may not be a viable solution as it is likely to decay and cause detrimental pollution. In the near future, automatic feeding systems that maintain steady concentrations of algae between water exchanges will help mitigate this possible limitation.

Our baseline diet comprises a simple combination of *Chaetoceros sp.* and *Isochrysis* sp. (T. iso). A diverse diet is thought to provide more complete nutrition and promote sexual maturation, even though animals cultured with a single food type can reach sexual maturity (Joly *et al*. 2007). Reasoning that a food supplement could improve the quality and quantity of egg production, we tested the Oyster feast and showed that it enhanced gamete production, at least when other foods were potentially limiting. As supplements also add calories, and excess food can cause pollution detrimental to early stages, we only added Oyster Feast after approximately 2 months, when most of the animals had orange pigment organs. Other supplements have been used in *Ciona* culture (Joly *et al*. 2007; Zupo *et al*. 2020) and warrant additional test using our protocol with smaller tank system and size selection.

In our previous culture system (Ohta *et al*. 2020),we successfully used the BioDigest (Prodibio) probiotic as microbial supplement with so-called “bioballs”, but omitted this addition in the current study (while we tested it in a part of our 4^th^ round of culture), where we compensated its absence by frequent water exchanges afforded by the lower volumes. Given the observed impact of excess food and pollution on water quality and animal growth and maturation, we anticipate that controlling for a healthy microbiome may improve both animal digestion and physiology, and water quality (Wei *et al*. 2020; Liberti *et al*. 2021).

Finally, general environmental parameters, which we did not assay, may also influence animal growth and maturation in culture. Chief among them, temperature is known to affect growth and maturation in *Ciona*. Developmental speed is typically correlated with temperature within their range of thermal tolerance (Therriault and Herborg 2008; Wilson *et al*. 2022). Higher temperatures accelerate growth and maturation, while just a few degrees of excessive heat damages embryonic development and could also impact post-embryonic stages (Caputi *et al*. 2015; Sato *et al*. 2015, 2022; Malfant *et al*. 2017; Irvine *et al*. 2019). *Ciona robusta* is more tolerant to higher temperatures and salinity than *Ciona intestinalis* (Satou *et al*. 2015; Malfant *et al*. 2017; Wilson *et al*. 2022), and could in principle close the life cycle faster above 20 °C, compared to the 14 ℃ and 18 ℃ used in our culture system.

Other environmental conditions, such as light quality, intensity and circadian oscillations, as well as seasonality may also affect animal cultures. Indeed, *Ciona* adults accumulates eggs when placed under constant light, and controlling the circadian rhythm can be used to control spawning (Joly *et al*. 2007; Veeman *et al*. 2011).

In summary, while our provisional protocol still has room for improvement on multiple fronts, our principled science-based approach to study the impact of culture conditions on animal growth and maturation has yielded novel insights into post-embryonic developmental physiology, and produced a simplified and accessible protocol to culture both *Ciona robusta* and *Ciona intestinalis* in laboratory conditions. Future development will further improve this protocol and use it to generate novel genetic reagents for a broad range of studies using ascidian models.

## Author contributions

B.T.M. performed maintenance, observation and measurement of animals in the 2nd, 3rd and 4th rounds of culture, and maintained algae.

M.O. performed maintenance, observation and measurement of animals in the 2nd and 3rd rounds of culture, and maintained algae with B.T.M..

B.P.D.M. performed maintenance, observation and measurement of animals in the 1st round of culture.

C.C. and V.P. performed maintenance, observation and measurement of animals in the 2nd and 3rd rounds of culture.

L.C. designed the study, acquired funding, oversaw work and progress and wrote the paper.

N.O. designed the study, oversaw work and progress, performed maintenance, observation and measurement of animals in the 2nd and 3rd rounds of culture, analyzed data, prepared figures and wrote the paper.

## Acknowledgments

We thank all the members of the Christiaen laboratory both in New York and Bergen for helpful comments for our culture, especially Keaton Schuster and Yelena Bernadskaya for help shipping *Ciona robusta* juveniles to Bergen from NY. This project has been supported by core funding from Michael Sars Centre, University of Bergen, to L.C. LC wishes to dedicate this work to the memory of Jean-Stéphane Joly, who was an early champion of *Ciona* cultures.

---

**Supplemental Figure S1.**
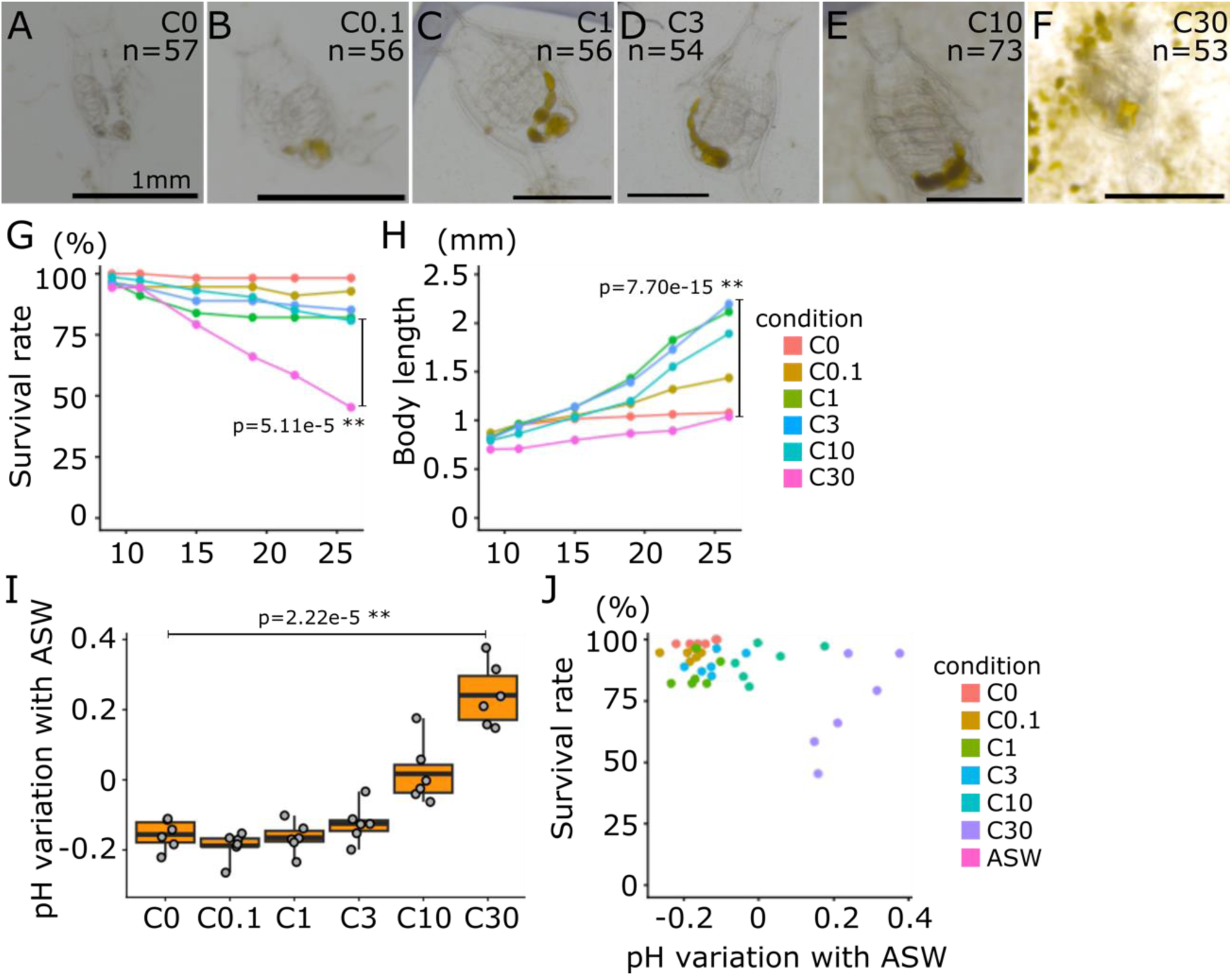
Survival and growth of *Ciona intestinalis* juveniles in Petri dish. (A-F) Images of juveniles at day 26 in each dietary condition are shown. bar=1mm (G) Survival rate of juveniles averaged in all plates in each dietary condition are shown. (H) Body length of individuals averaged in all plates in each dietary condition are shown. (I) pH in each dish is averaged in each dietary condition and standard with pH of artificial sea water (ASW) which was used for animal culture. (J) Scatter plot shows correlation between pH variation and the survival rate in each Petri dish in each dietary condition. Each dot shows individual condition which pH in the Petri dish was measured after culturing and averaged in each condition. Numbers of juveniles in each dietary condition are shown in A-F. The color indicates dietary conditions of the Petri dishes on G, H and J. *Ciona intestinalis* was used for all the graphs in this figure. p-value was calculated by Fisheŕs exact test in G and t-test in Hand I. p>0.05; N.S, 0.05>p>0.01; *, 0.01>p; **.

**Supplemental Figure S2.**
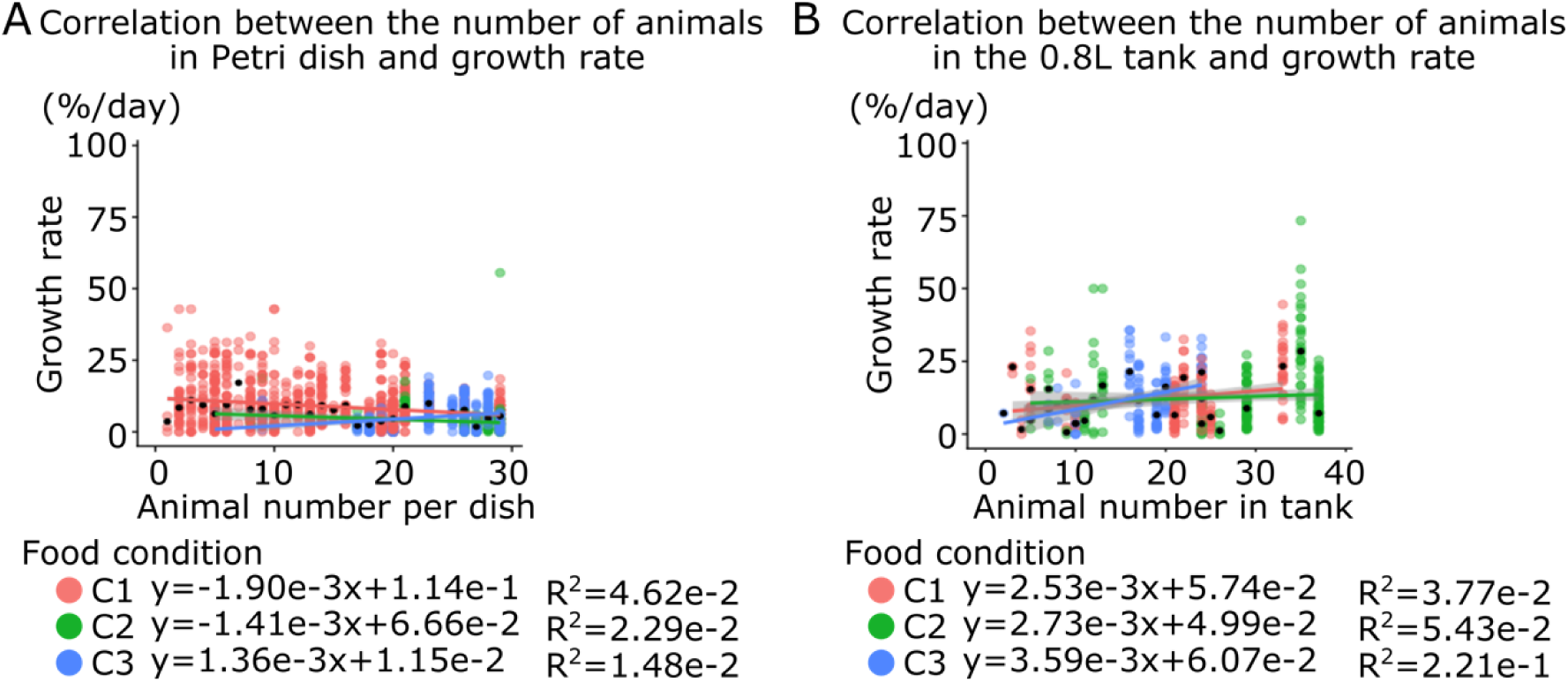
Correlation between density of animals and growth. (A) Correlation between density of animals in one Petri dish and growth rate is shown in scatter plot. (B) Correlation between density of animals in one the 0.8L tank and growth rate is shown in scatter plot. Each colored line indicates a fitting curve on A-B. Each dot shows individual animals observed at one time point on A-B. Black dots show median of the growth rate in each dietary condition on particle base or each condition of different animal numbers in a Petri dish on A-B. The color indicates dietary conditions of the Petri dishes on A-B. *Ciona robusta* was used for all the graphs in this figure.

**Supplemental Figure S3.**
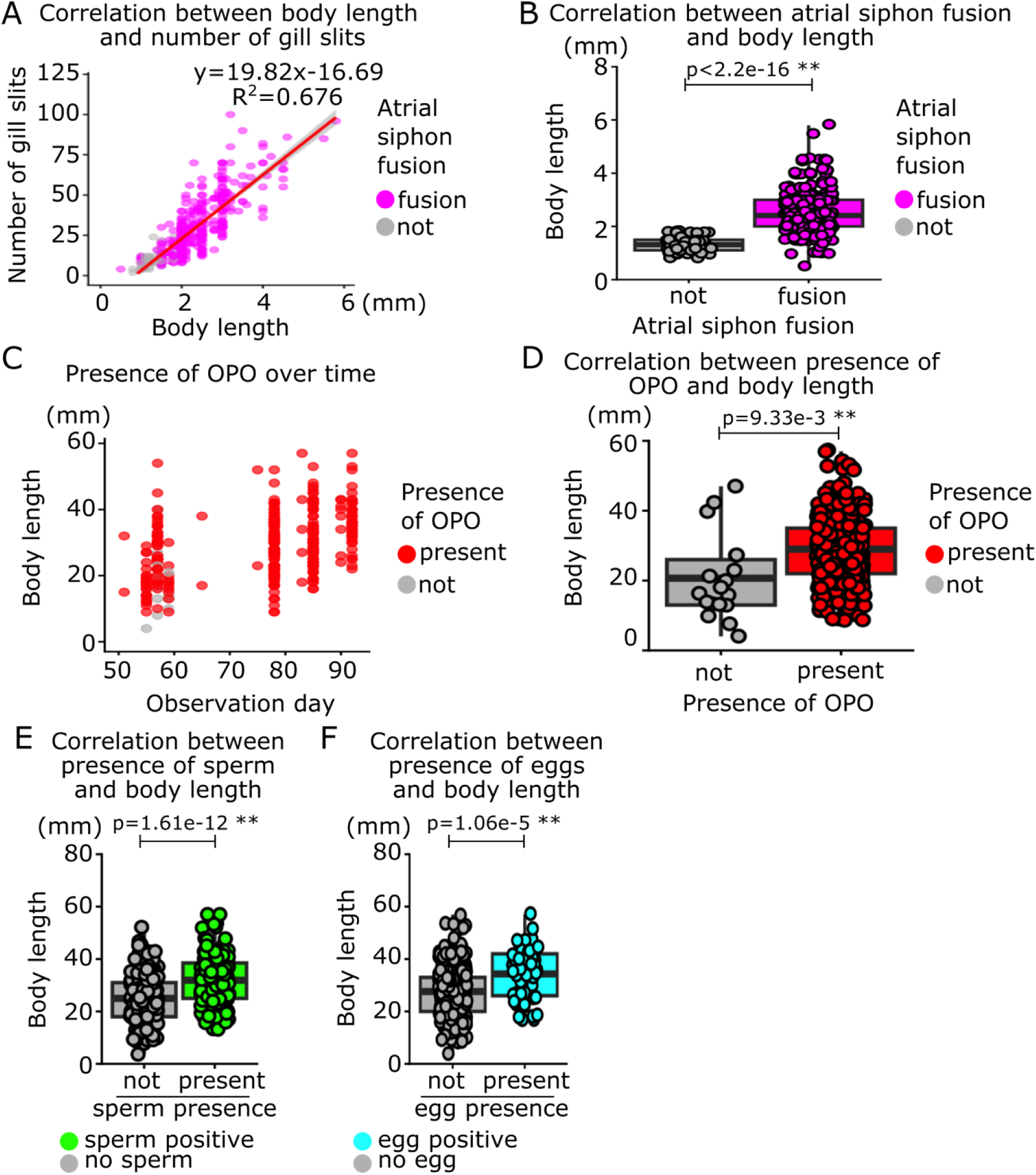
Growth and maturation of *Ciona robusta* in our 3^rd^ round of culture. (A) Correlation between body length, the number of gill slits and atrial siphon fusion is shown in scatter plot. The red line indicates a fitting curve. (B) Box and dot plot show correlation between atrial siphon fusion and body length. (C) Body length and presence of orange pigment organ (OPO) over time are shown in scatter plot. (D) Box and dot plot show correlation between presence of OPO and body length. (E) Box and dot plots show correlation between presence of sperm and body length. (F) Box and dot plots show correlation between presence of eggs and body length. Each dot shows individual animal observed at one time point on A-F. Magenta and gray show completion of atrial siphon fusion or not on A-B. Red and gray show presence of OPO or not at the tip of sperm duct on C-D. Green and gray show presence of sperm or not on E. Cyan and gray show presence of egg or not on F. *Ciona robusta* is used for all the graphs in this figure. p-value was calculated by t-test on B, D, E and F. p>0.05; N.S, 0.05>p>0.01; *, 0.01>p; **.

## Notes

### Competing Interest Statement

The authors have declared no competing interest.

